# Selection for protein stability enriches for epistatic interactions

**DOI:** 10.1101/338004

**Authors:** Anna Posfai, Juannan Zhou, Joshua B. Plotkin, Justin B. Kinney, David M. McCandlish

## Abstract

A now classical argument for the marginal thermodynamic stability of proteins explains the distribution of observed protein stabilities as a consequence of an entropic pull in protein sequence space. In particular, most sequences that are sufficiently stable to fold will have stabilities near the folding threshold. Here we extend this argument to consider its predictions for epistatic interactions for the effects of mutations on the free energy of folding. Although there is abundant evidence to indicate that the effects of mutations on the free energy of folding are nearly additive and conserved over evolutionary time, we show that these observations are compatible with the hypothesis that a non-additive contribution to the folding free energy is essential for observed proteins to maintain their native structure. In particular through both simulations and analytical results, we show that even very small departures from additivity are sufficient to drive this effect.

## 1. Introduction

The relationship between protein sequence, stability, and function has been a subject of intense investigation for decades. A combination of biophysical and evolutionary models and, more recently, high-throughput mutagenesis experiments have dramatically advanced our understanding of this complex relationship [1–5]. A consensus view has emerged on some aspects of protein functions and evolution— e.g., what accounts for the distribution of thermodynamic stabilities observed in nature. And yet other questions—e.g., whether genetic interactions play a dominant or minor role in protein sequence evolution—remain actively debated, with apparently contradictory empirical and theoretical evidence [1–5].

A nuanced appreciation of the high-dimensional nature of protein sequence space has been essential for resolving questions about protein structure, function, and evolution. The observation that naturally occurring proteins are only marginally, as opposed to maximally, stable was first interpreted as an adaptive feature to permit increased protein flexibility and functionality [6]. But, with some exceptions [7], this view has been largely replaced with a more parsimonious explanation based on the high dimensionality of sequence space: marginal stabilities are observed because, simply, far more sequences are marginally stable than maximally stable [8]. Essential to the development of this explanation was the concept of sequence entropy [9,10] – the idea that the sheer number of protein sequences that map to a given phenotype exerts a strong entropic pull on the distribution of observed phenotypes in viable proteins [11–13]. The field today has mostly settled on a synthetic understanding of how simple biophysical models of energy and folding, along with the structure of sequence space, conspire to explain the distribution of protein stabilities observed in nature [1–5].

By contrast to observed stabilities, the role of epistasis in protein evolution and function remains a topic of active debate with unresolved ambiguities. The same biophysical models that can parsimoniously explain observed distributions of stabilities have been reported to show only a weak context-dependence of mutational effects on stability [14], or, alternatively, reported to show very strong context dependence of mutational effects [13,15,16]. Likewise, experimental studies on the fitness effects of mutations in divergent sequence backgrounds have reportedly very weak epistasis [14], whereas comparative analysis of divergent proteins has implicated an overriding role for epistasis in shaping sequence evolution [17]. How are we to resolve this significant discrepancy about the role of epistasis for protein stability and sequence evolution?

In this paper we address this discrepancy by analyzing simple models for the relationship between amino acid sequence and the Gibbs free energy (Δ*G*) of folding. Under selection to maintain a minimum degree of stability, these models predict distributions of folding energies that are roughly consistent with those observed in nature. Moreover, the models predict very weak interactions between pairs of mutations. These predictions are consistent with biophysical measurements of nearly additive mutational effects on stability [18], and with reports of consistent effects over both short [14] and long [19] evolutionary timescales. And yet, at the same time, we show that a non-additive contribution to the folding free energy is essential for allowing proteins to fold stably in our model, for reasons attributable to sequence entropy. These results may help to resolve striking discrepancies in the literature on the importance of epistasis for protein stability and evolution [1–5].

## 2. Methods

### 2.1. Simulations

We consider two simple models for the relationship between amino acid sequences and ΔG of folding. All amino acid sequences considered are of length l = 400. Both of these models have the form:

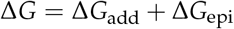

where Δ*G*_add_ describes the independent and additive contributions from each amino acid at each position, while Δ*G*_epi_ describes contribution of interactions between positions.

First we consider a model where epistasis arises solely from pairwise interactions between sites. For example, Δ*G*_add_ in such a model might account for the independent energetic effects of individual residues being removed from the aqueous environment upon protein folding, whereas Δ*G*_epi_ might account for physical interactions between pairs of residues in the folded conformation. For this model, we assume that the additive effect on stability for each possible amino acid in each position in the primary sequence is drawn from a Gaussian distribution with mean *μ_add_* and variance 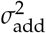. In addition, we allow pairwise interactions between amino acid sites, with the magnitudes of these interactions drawn from a Gaussian distribution with zero mean and variance 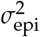. Moreover, these pairwise interactions are specified in such a way that the pairwise interaction terms have no impact on the average effect of any given amino acid substitution, so that the magnitudes of the additive and epistatic effects can be modified independently. That is, the model is equivalent to a “random field model” from the fitness landscape literature [20,21] where the only non-zero terms are the constant, linear, and pairwise interaction terms. See Appendix A for details on the mathematical features and practical implementation of this model.

Second, we consider a model where epistasis is modeled as a random deviation from additivity drawn independently for each *genotype*, meaning each sequence of amino acids. This model is similar to the “rough Mount Fuji” model of fitness landscapes [22,23], and might arise if Δ*G*_epi_ accounts for highly cooperative energetic contributions, e.g., resulting from global changes in the folded protein conformation. In this case we again draw the additive effect of each amino acid in each position from a Gaussian distribution with mean μ_add_ and variance 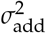, but in addition the folding energy of each genotype is perturbed by an independent draw from a zero-mean Gaussian with variance 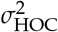 (where *HOC* denotes “house of cards”, since this component is completely uncorrelated between mutationally adjacent genotypes, similar to the house of cards model of fitness landscapes [24]). Because protein sequence space is too large to store in computer memory, we implement a hashing scheme so that in the simulations these epistatic effects remain consistent for previously observed genotypes, but are drawn anew for genotypes that have not yet been encountered.

The simulations of protein sequence evolution under selection are based on a threshold model for thermodynamic stability: proteins with a negative Δ*G* of folding are deemed viable and all other sequences are deemed inviable. At each step in the simulation, a random position in the protein sequence is chosen and changed to a random alternative amino acid. This new sequence is accepted if it is viable and rejected if it is inviable; if the mutation is rejected then the protein sequence remains unaltered for that step of the simulation. These simulations are initialized at the sequence predicted to be most stable based on its additive effects and allowed to equilibrate for 5000 proposed mutations, a time sufficient for the distribution of folding stabilities to become approximately stationary for the conditions considered. After this relaxation period, the simulations continue for an additional 5000 proposed mutations to produce the results shown here. All simulations and all calculations presented were implemented in Mathematica and the corresponding Mathematica notebook is included as supplemental information.

The models analyzed here are simpler and less realistic than other commonly used models for protein evolution based on force fields [25], contact energies [26], or lattice proteins [27]. However, we employ these models because their simple structure facilitates a variety of exact and approximate analytical results, and thus provides a clearer illumination of the theoretical issues involved than the more realistic but less tractable alternatives.

## 3. Results

### 3.1. Epistasis is essential for proper folding of evolved sequences

We simulated the evolution of a protein of length 400 under a model where each amino acid at each position makes an additive contribution to the free energy of folding, and where in addition we allow pairwise stability interactions between sites. We imposed truncation selection for spontaneous folding so that only sequences with a negative Δ*G* of folding are considered viable. The parameters of the simulations were chosen to be roughly consistent with the observed distribution of folding stabilities and mutational effects on stability reported in the literature [e.g. 28-31]

Figure 1a shows the distribution of folding energies observed in these simulations after the process was allowed to reach stationarity. The mean of this distribution is only slightly negative, indicating that the evolved proteins are marginally stable, as predicted by theory and observed in nature [8,11,29,31,32]. Examining the effects of random single amino acid substitutions for sequences drawn from this distribution (Figure 1b), we observe that the mean effect is +1.15 kcal/mol with standard deviation 1.5 kcal/mol, consistent with empirical observations [28–30]. The distribution of energetic effects of mutations that are fixed over the course of the simulations (Figure 1c) is shifted to have approximately zero mean (−0.0007 kcal/mol) and a smaller standard deviation (1.1 kcal/mol) as observed previously [16,33,34].

**Figure 1.**
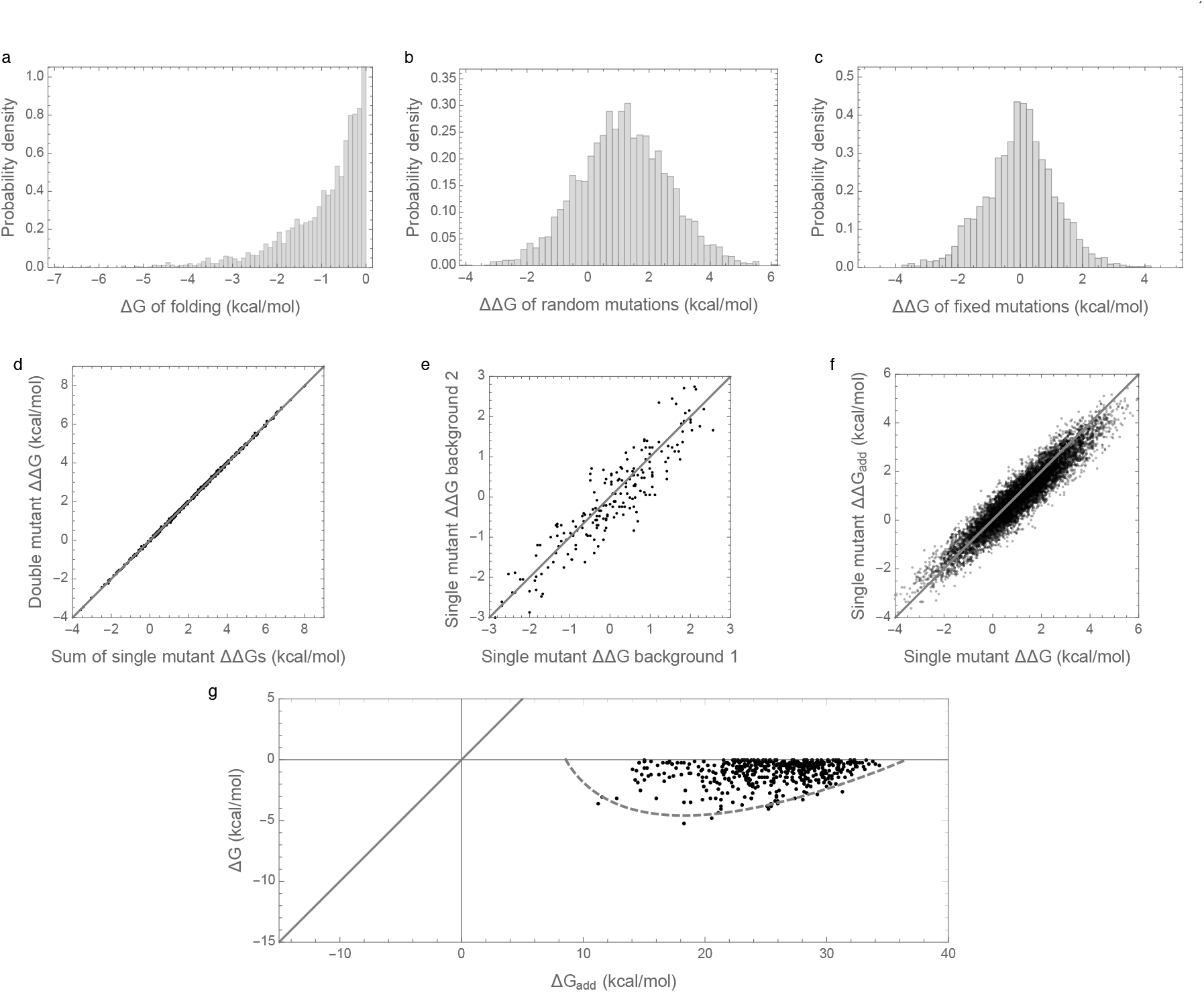
Free energy of folding, stability effects of mutations, and contribution of additive effects to folding stability under the pairwise epistatatic effects model: (**a**) Distribution of free energy of folding for evolved sequences. (**b**) Distribution of stability effects of random mutations, i.e. distribution of ΔΔ*G* for a random single mutant generated from a random evolved sequence. (**c**) Distribution of stability effects of fixed mutations, i.e. distribution of ΔΔ*G*s corresponding to two distinct neighboring sequences along the simulated trajectory. (**d**) Stability effects of double mutations versus the sum of the stability effects of the two single mutations. 500 random double mutants are shown, *R*^2^ = 0.9997. (**e**) Effects of single mutations that fixed along the trajectory in two evolved backgrounds that differ by 50% sequence divergence, *R*^2^ = 0.87. (**f**) The stability effect of a random mutation on Δ*G* is highly correlated with the stability effect of the mutation on the additive contribution to free energy Δ*G*_add_, *R*^2^ = 0.89. (**g**) Free energy of folding versus additive contribution to free energy of folding for evolved sequences. The additive contribution to folding is not a good indicator of the total free energy of folding (*R*^2^ = 0.06) and observed sequences cannot fold spontaneously based on the additive contribution alone. The solid curve is derived from our bivariate normal approximation and is predicted to contain 95% of the evolved sequences. Simulations conducted under the pairwise epistasis model with *μ*_add_ = 1 kcal/mol, 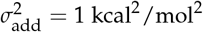, and 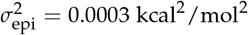.

Interactions between mutations also have a similar magnitude to those observed in previous studies. In an evolved background, the combined mutational effect of two random mutations is nearly exactly predicted by the sum of the mutational effects of the constituent single mutations (Figure 1d, *R*^2^ = 0.9997). Furthermore, the effects of mutations that fix along our simulated evolutionary trajectories remain relatively consistent over time (Figure 1e), with a root mean square change of only 0.5 kcal/mol at 50% sequence divergence, consistent with the empirical measurements of Risso *et al*. [19], who observed an RMS change of .67 kcal/mol among mutations that have fixed between two sequences at a similar level of divergence. Because we are working with simulated data, we can also assess how closely the observed effects of mutations reflect the effects of these mutations on the additive component of the folding energy. Figure 1f shows that the observed effects of random single amino acid substitutions, i.e. the ΔΔ*G* of single amino acid substitutions, are highly correlated with the underlying additive effects of these mutations, i.e. with the ΔΔ*G*_add_ of the same mutations (*R*^2^ = 0.89).

To summarize, our simulations are qualitatively similar to both previous empirical and theoretical investigations of long-term evolution under selection for protein folding stability, and, on the face of it, based on Figures 1d, 1e, and 1f, they suggest that epistasis plays only a minor role in protein folding. However, when we actually compute the additive contribution to protein stability observed in our simulations, a very different picture emerges as shown in Figure 1g. Shockingly, we find that the additive contribution to folding stability is not nearly sufficient to allow spontaneous folding (mean of Δ*G*_add_ is 25.5 kcal/mol), and so epistatic interactions are required for folding of all sequences observed at stationarity. We also observe that the additive contribution to folding stability Δ*G*_add_ is almost completely uncorrelated with the actual folding stability Δ*G*, with *R*^2^=0.06. Thus, in our model, epistasis plays an essential role in protein folding, despite the fact that mutational effects can be well predicted based solely on the additive contributions of each amino acid to the sequence’s free energy of folding.

### 3.2. Enrichment for epistasis observed under pairwise, but not independent model

In order to better understand the causes of these counter-intuitive results, we considered an alternative model with an identical additive component but with epistasis modeled as an independent random draw for each sequence (from a zero-mean Gaussian distribution with variance 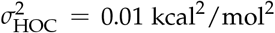, see Methods). The results of these simulations are shown in Figure 2. The distribution of folding stabilities, the distribution of mutational effects, and the extent of observed epistasis observed in mutational effects are qualitatively unchanged from the results observed under the prior, pairwise interaction model (Figure 2a-f vs. Figure 1a-f). However, in this case the paradoxical contribution of epistasis to folding stability is absent, so that the additive contribution to stability is sufficient for spontaneous folding for most evolved sequences, and the observed folding energy is highly correlated with the additive contribution (*R*^2^=0.99, Figure 2d). We therefore conclude that enrichment for epistasis under stabilizing selection occurs with pairwise epistatic interactions but not with fully random interactions.

**Figure 2.**
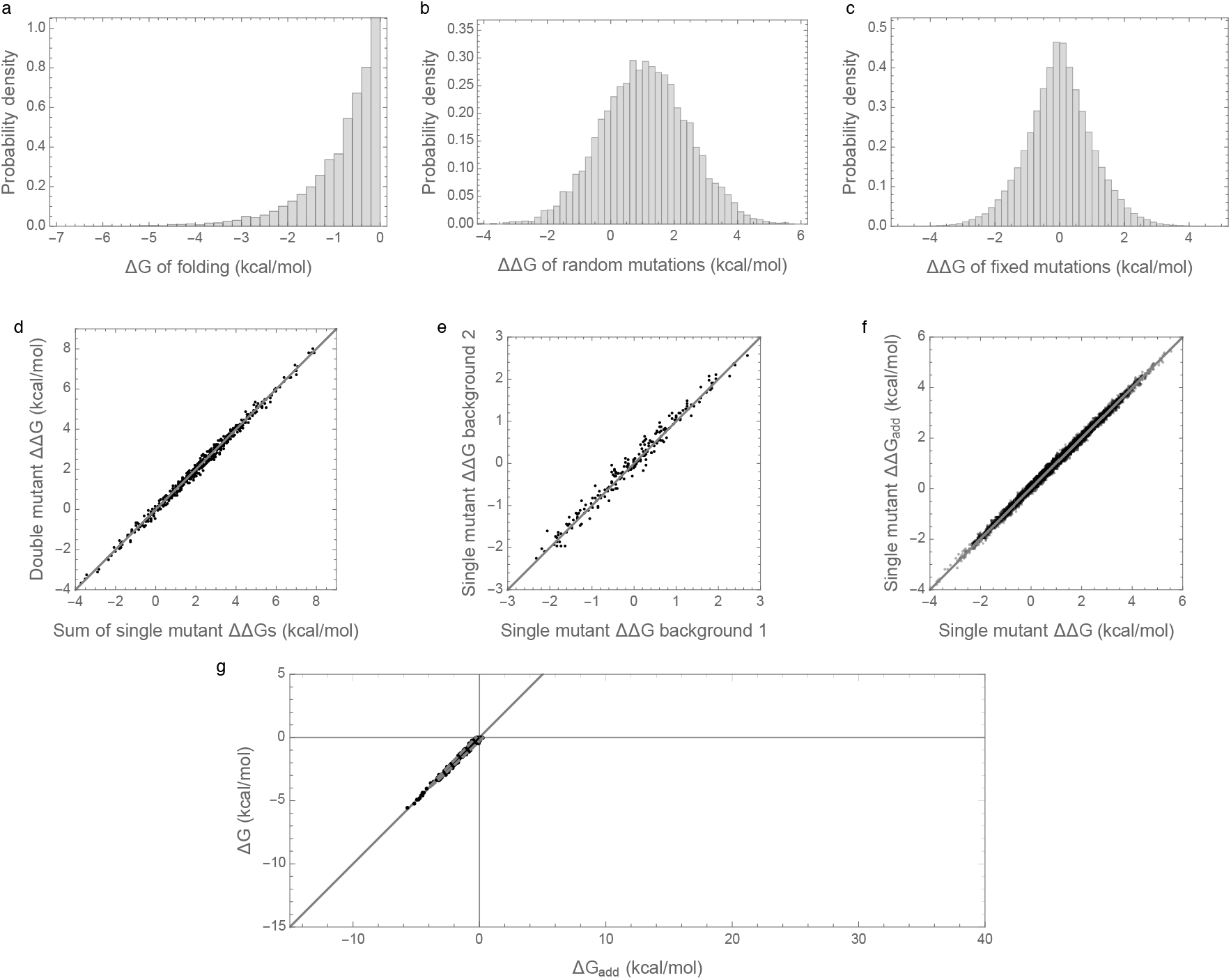
Free energy of folding, stability effects of mutations, and contribution of additive effects to folding stability under the independent epistatic effects model: (**a**) Distribution of free energy of folding for evolved sequences. (**b**) Distribution of stability effects of random mutations, i.e. distribution of ΔΔ*G* for a random single mutant generated from a random evolved sequence. (**c**) Distribution of stability effects of fixed mutations, i.e. distribution of ΔΔ*G*s corresponding to two distinct neighboring sequences along the simulated trajectory. (**d**) Stability effects of double mutations versus the sum of the stability effects of the two single mutations. 500 random double mutants are shown, *R*^2^ = 0.993. (**e**) Effects of single mutations that fixed along the trajectory in two evolved backgrounds that differ by 50% sequence divergence, *R*^2^ = 0.98. (**f**) The stability effect of a random mutation on Δ*G* is highly correlated with the stability effect of the mutation on the additive contribution to free energy Δ*G*_add_, *R*^2^ = 0.997. (**g**) The additive contribution to folding is a good indicator of the total free energy of folding (*R*^2^ = 0.99) and 95% of observed sequences can fold spontaneously based on the additive contribution alone. The dashed gray curve is derived from our bivariate normal approximation and is predicted to contain 95% of the evolved sequences. Simulations conducted under the independent epistatic effects model with *μ*_add_ = 1 kcal/mol, 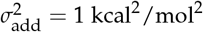, and 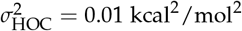.

What explains this difference in behavior between the model with pairwise epistatic interactions and the model with independent epistatic effects for each sequence? In order to address this question, we conducted a mathematical analysis of the random field model with pairwise epistatic interactions (see Appendix B). What we came to understand was that, in this model, the amount of epistasis observed in double mutants vastly underestimates the total amount of epistasis in the energy landscape. This occurs because making a double mutant only results in changes to relatively few interaction pairs (i.e. those interaction pairs involving the site of either single mutant). However, as additional mutations are added to the sequence, more pairs are perturbed, which unleashes additional epistasis.

More precisely, in the mathematical analysis we considered the expected magnitude of the observed epistasis as a function of the number of mutations from an arbitrary focal sequence. That is, we calculated the expected variance in the epistatic contribution among the set of all sequences at a given distance *d* from the focal sequence. The results are shown in Figure 3 where the variance at *d* = 2 is set to 1, so that the variance is expressed relative to the variance observed in a double mutant analysis. We see that for small *d* this variance increases roughly linearly, and eventually saturates at almost 50 times the expected variance at *d* = 2. In contrast, the independent random epistasis model is essentially constant at all positive distances. Thus, similar levels of observed epistasis in double mutants make vastly different predictions for the total amount of epistasis under the two models.

**Figure 3.**
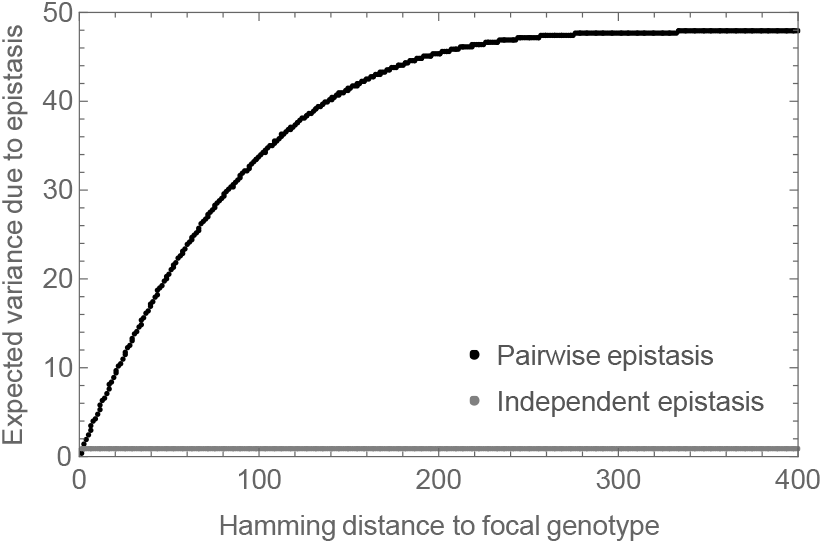
Expected epistatic variance as a function of distance from the focal sequence for amino acid sequences of length *l* = 400. Results for the pairwise model shown in black, results for the independent epistasis model shown in gray. All variances are normalized relative to the expected variance at *d* = 2 which is set to 1. Notice that the epistatic variance at large distances is much larger than the epistatic variance at distance *d* = 2 for the pairwise epistasis model but not for the independent epistasis model.

### 3.3. Bivariate normal approximation for joint distribution of additive and epistatic contributions to the free energy of folding captures impact of sequence entropy

Our results on the surprising implications of small observed epistatic effects under the pairwise interaction model make the results in Figure 1g appear somewhat more plausible because more epistasis is present in the landscape than is apparent from the double mutants. But this observation still does not provide a definite explanation for the large contribution of epistatic interactions to folding stability.

We now provide such an explanation, based on considering the fraction of random sequences that have any given pair of additive and epistatic contributions (see Appendix C for details). In particular, we assume that the distribution of additive contributions to the free energy of folding for random sequences is normally distributed with mean *μ*_1_ = *lμ*_add_ and variance 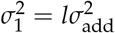. We further assume that the distribution of epistatic contributions is normally distributed with mean 0 and variance 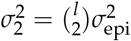. The additive and epistatic contributions are uncorrelated, so the total folding energy of a random sequence ΔG = Δ*G*_add_ + Δ*G*_epi_ is normally distributed, with mean *μ* = *μ*_1_ and variance 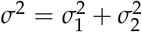. These normal approximations are reasonable considering that ΔG_add_ is calculated by adding up the energy contribution of each site in the sequence, and ΔG_epi_ is calculated by adding up the energy contribution of each pair of sites in the sequence, for a large number *l* = 400 of sites. We also note that in the mutation-limited regime depicted in our simulations, the stationary distribution of the simulated random walk will be uniform on the set of genotypes with negative folding energies that are path-connected with our choice of starting genotype [35]. Under the assumption that almost all genotypes with negative folding energies are path-connected, picking a sequence from the stationary distribution is equivalent to picking a sequence from the uniform distribution on sequences with negative folding energies, and so our problem reduces to understanding the distribution of additive and epistatic folding contributions among all sequences with negative free energies of folding.

Under the above approximation, we now consider how—for a typical viable sequence— the additive and epistatic energies jointly produce a negative free energy of folding. The key idea is that there are so many more sequences with positive additive contributions to folding than there are sequences with negative additive contributions that most sequences that fold have a positive additive contribution despite the fact that any particular sequence with a positive additive contribution to the free energy of folding has only a minuscule chance of actually folding.

More precisely, let us fix the value of the additive energy at ΔG_add_ = *x*, and count the number of sequences, with this given ΔG_add_, that fold. The number of sequences with ΔG_add_ = *x* is proportional to the probability density for the distribution of ΔG_add_, 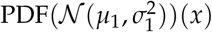. Adding the epistatic energy to the additive energy, the sequences with ΔG_add_ = *x* that fold are the sequences for which ΔG_epi_ < −*x*, i.e. their number is proportional to the cumulative distribution function of Δ*G*_epi_ evaluated at −*x*, 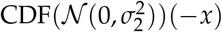. Putting the two pieces together, the number of sequences that have ΔG_add_ = *x* and, at the same time fold, is proportional to

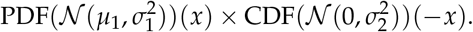

Figure 4 shows this calculation for *x* values near the viability threshold 0. We see that over this range of folding energies the number of sequences is growing extremely rapidly (Figure 4a) so that, roughly speaking, the number of sequences with a given additive energy increases 10-fold for every additional .45 kcal/mol. Now, Figure 4b shows the fraction of sequences with a given additive contribution that spontaneously fold. This is near 1 for most sequences with a negative contribution, but decreases exponentially for positive additive contributions. The net result (Figure 4c) is that a typical additive contribution for a sequence that folds is often around 22 or 23 kcal/mol. While only a tiny fraction of sequences with additive energies in this range fold (roughly 1 in a million), there are roughly 10 billion times more sequences in this 1 kcal/mol range than there are all sequences that would spontaneously fold based on their additive contribution (i.e. with ΔG_add_ < 0), so that in the end most sequences that fold have substantially positive additive contributions to the free energy of folding.

**Figure 4.**
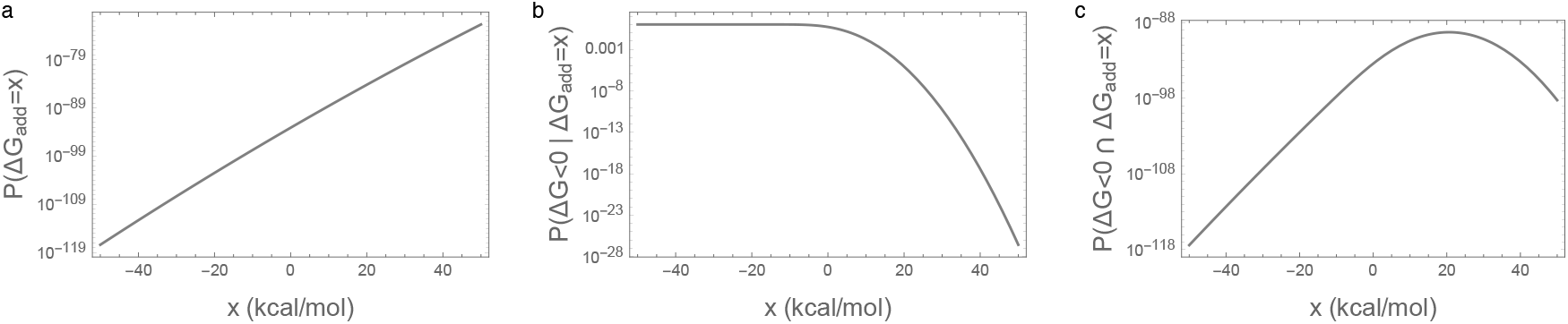
Illustration of the main mechanism behind the essentiality of epistatic interactions for spontaneous folding: (**a**) Density of random sequences with given additive free energy 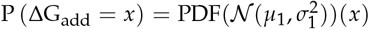. (**b**) Fraction of sequences that fold given additive free energy 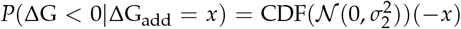. (**c**) Density of random sequences that fold and have the given additive free energy P (ΔG < 0 ∩ ΔG_add_ = *x*). Parameters are identical to those in Figure 1.

The above argument leads to a simple prediction for the joint distribution of the free energy of folding and additive contribution to folding shown in Figures 1g and 2g: since the joint distribution for random sequences is bivariate normal, the distribution of observed energies should simply be this bivariate normal distribution truncated at ΔG = 0 kcal/mol. This approximation is shown in Figures 1g and 2g by a dashed gray curve that is predicted to contain 95% of the observations, and we see that this prediction is in reasonable agreement with our simulations.

Moreover, under this bivariate normal approximation, the average contribution of epistasis to the mean free energy of folding observed in our simulations can be calculated in a manner exactly analogous to Galton’s classical results on regression to the mean [36], or the difference between the selection differential and the response to selection in the breeder’s equation from quantitative genetics [37,38]. In particular, we find that the mean additive energy of viable sequences is approximately 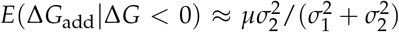 (see Appendix C for details), so that the mean contribution of epistasis is approximately 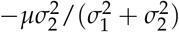, or equivalently −*μ*(1 − *R*^2^), where *R*^2^ is given by the squared correlation coefficient of additive and total folding energies taken over all of sequence space. As a result, even if the mapping from sequence to folding energy is nearly additive, in the sense that *R*^2^ is almost 1, the predicted epistatic contribution to the folding stability can still be substantial provided that the expected folding energy *μ* of a random sequence is sufficiently disfavorable.

## 4. Discussion

The role of epistasis in long-term protein evolution remains a topic of active debate [1–5]. Here we have explored a surprising phenomenon where the effects of mutations on the Δ*G* of folding appear to combine nearly additively, and nonetheless what little epistasis is present plays a critical role, to the extent that observed sequences would not be able to fold in the absence of these epistatic interactions. We showed that this phenomenon occurs in a model where interactions occur between pairs of sites but not in a model where each sequence differs from its additive prediction by an independent draw from a normal distribution. The difference between the two models arises because pairwise interactions can appear nearly additive in double mutants while still producing a substantial amount of epistasis over sequence space as a whole. We also present simple analytical approximations that predict the extent of the epistatic contribution to stability in our simulations. Furthermore, these approximations suggest that this phenomenon occurs due to sequence entropy: many more sequences can fold due to a combination of epistatic and additive contributions than can fold based on the additive contributions to stability alone, and so the epistatic contribution to stability is typically essential when one observes a random sequence that folds. These results add to a growing literature demonstrating that natural selection can enrich for epistatic interactions in both adaptive [39–41] and nearly neutral [13,16] evolution, such that the mutations that fix during evolution can have a very different pattern of epistasis than random mutations.

Our simulations (Figure 1) recapitulate the known qualitative features of protein evolution under purifying selection for folding stability to a surprising degree, with the exception of matching the observed stability margin, which is smaller in our simulations than for experimentally measured folding energies [29] (free energy of folding is typically −5 to −10 kcal/mol versus −1 kcal/mol in our simulations). However, this unnaturally small stability margin is a well-known artifact of our decision to model fitness as a step function in stability [8] rather than a more realistic logistic function [11,30], and the fact that our simulations do not include any of the other factors that would tend to increase the stability margin such as selection for mutational robustness [8,32,35] or selection to prevent misfolding due to errors in translation [42]. Nonetheless the simple sequence-to-fitness mapping employed in our simulations allows us to provide a relatively simple and complete theory for the observed phenomenon. Moreover, we emphasize that it is easy to find realistic parameters where the mean additive contribution to stability is far less stable than shown in Figure 1, so we anticipate that the possibility that most sequences fold only due to epistasis would be robust even if sequences experienced a much larger stability margin.

A different limitation of our results concerns the assumption, in our truncated bivariate normal approximation, that the set of sequences with negative folding energies is mutationally connected, and hence accessible to an evolving population. In particular, the theory breaks down if a large fraction of sequences that fold appear as isolated peaks or small isolated clusters of sequences. Figure 5 shows an example of this limitation for the case of the independent model with parameters chosen so that the bivariate normal approximation is identical to the bivariate normal approximation for the pairwise model shown in Figure 1. The figure shows some enrichment for epistasis but not as much as predicted by our bivariate normal approximation. Using the crude percolation-theory argument that the connected network of sequences can extend only up to the additive energy at which each sequence has on average one neighbor that folds due to epistasis [43], we can derive the approximate upper limit of the distribution of additive energies as −*σ*_HOC_**Ψ**^−1^ (1/ (400 × 19)) = 18.8, where **Ψ**^−1^ is the inverse cumulative distribution function of a standard normal distribution. This approximate upper limit is shown by the dashed vertical line in Figure 5. We see that the cloud of observed sequences is primarily to the left of this line, with a notable absence of sequences with substantially more positive additive contributions. This analysis of connectivity of the set of sequences that fold highlights that pairwise interactions have several special features: not only can they appear locally non-epistatic while harboring a substantial amount of epistasis at greater distances, but as long as the individual coefficients remain small they produce energy landscapes that change smoothly over sequence space, producing the enormous connected networks of sequences whose traversal allows the evolution of a sizable epistatic contribution to folding.

**Figure 5.**
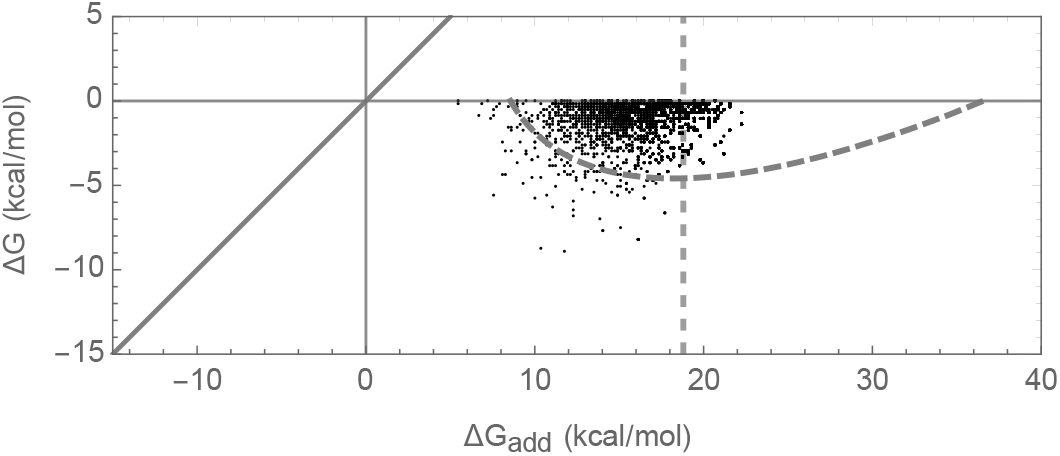
Joint distribution of Δ*G* of folding and the additive contribution to Δ*G* of folding for the independent epistasis model with 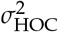 chosen so that the bivariate normal approximation matches the bivariate normal approximation shown in Figure 1g. Simulations conducted under the independent epistasis model with *μ*_add_ = 1 kcal/mol, 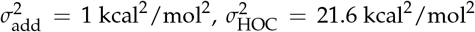. Dashed curve shows area predicted to include 95% of sequences at stationarity under the bivariate normal approximation; dashed vertical line shows approximate left-most edge of region where bivariate normal approximation is valid based on a crude percolation theory argument (see text).

A final limitation of our approach concerns long-term dynamics at individual sites. Our analysis addresses the possibility that the observed, nearly additive effects of mutations on stability may be compatible with a critical role for epistasis in most sequences that fold. But our mathematical results do not yet provide a detailed view of how the stability effects at individual sites change over time. In particular, previous simulation studies have described complex temporal dynamics at individual sites, so that the set of tolerable amino acid mutations at any given site changes over time [13,15,16,44,45], a process that has been called an “evolutionary Stokes shift”, and which is also related to the phenomena of “contingency and entrenchment” and the expectation that reversion rates will decrease over time [15,45–47]. However, mathematical results closely related to the approach taken here [13,48] suggest that extensions of our theory to address the temporal variation in stability effects, as well as the corresponding distributions of site-specific amino acid preferences and correlations between sites, may also be possible.

It is natural to ask whether the essentiality of epistatic interactions for the functionality of evolved sequences is likely to hold in other contexts where additivity is thought to prevail, such as the DNA binding sites of transcription factors (TFs). However, this effect is unlikely to occur in such cases because the sequences are much shorter, the alphabet is smaller, and thus the set of functional sequences makes up a much larger proportion of genotypic space. In particular, it is helpful to consider the z-score of functional sequences relative to random sequences, since the regression to the mean effect observed here is proportional to the absolute value of the z-score. For instance, a typical TF binding motif in bacteria has an information content of ~ 23 bits [49], corresponding to a p-value of ~ 10^−7^ or a z-score of roughly −5. Eukaryotic TFs have even smaller information content and therefore smaller absolute value z-scores. In contrast, the z-scores of the spontaneously folding sequences observed in our simulations are on the order of −20, which we would expect to result in a roughly 4-fold larger contribution of epistasis to binding energy at stationarity than for a bacterial transcription factor binding site. Such extreme z-scores are not even possible in short DNA elements, e.g., the most extreme z-score possible in a TF binding site of length 20 is only −7. Thus, the essentiality of epistatic interactions observed here is likely possible only because protein sequence space is very large compared to the space of TF binding sites.

Finally it is important to emphasize that the key question of whether epistatic interactions for protein stability are essential for protein folding in naturally evolved sequences remains open. Our contribution only demonstrates that such an effect is qualitatively consistent with empirical observations on the thermodynamic effects of mutations and the results of prior simulation studies, and suggests that the overall importance of epistasis for stability depends on the precise form of epistasis involved. Intriguingly, the experimental observation that pairwise correlations between site-specific amino acid usages are sometimes necessary for folding [50] provides evidence for both the presence of the low-order epistatic interactions that result in a substantial contribution of epistasis to protein folding and also for the possible essentiality of these interactions. Thus, determining whether epistasis is essential for folding of observed sequences is a key question for the field, from both theoretical and empirical perspectives. Importantly, our analysis shows that most standard designs for examining the extent of epistasis for protein stability cannot adjudicate this question, because they examine how the stability effects of mutations change at a only single distance from a reference genotype. For instance, the analysis of double mutants considers the change in the effect of a mutation in a sequence at distance 1; and comparison of the effects of mutations on two diverged backgrounds, e.g. [14,19], can only determine the extent of epistasis at that one level of divergence. Rather, the two theories analyzed here differ in how the extent of epistasis changes with distance (e.g. Figure 3 and Appendix B.1). Thus, the critical experiment is to measure how the energetic effects of individual mutations change across several different levels of sequence divergence (c.f. [16]).

## Author Contributions

Conceptualization, A.P., J.B.P, J.B.K., and D.M.M.; Software, A.P., J.Z., and D.M.M.; Formal Analysis A.P., J.Z., J.B.K. and D.M.M.; Investigation A.P. and D.M.M.; Writing - Original Draft Preparation, A.P., J.Z., J.B.P, J.B.K., and D.M.M.; Writing - Review & Editing, A.P., J.B.P., J.B.K., and D.M.M.; Funding Acquisition, J.B.P and J.B.K.

## Acknowledgments

We thank Ashley Teufel and David Liberles for organizing this special issue of Genes and two anonymous reviewers for helpful comments.

## Appendix A

### Appendix A.1 Model for the free energy of folding

Given an alphabet 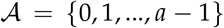 and a sequence length l, let 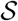 be the set all possible strings of length *l* built from alphabet 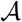. The free energy of folding ΔG(*x*) for a sequence 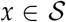 is defined as the sum of (1) an additive component that measures the energy contribution of each allele at each position in the sequence, and (2) an epistatic component that describes the energy contribution of pairwise interactions among alleles for the pairwise model, or a random draw from a normal distribution for the independent epistasis model:

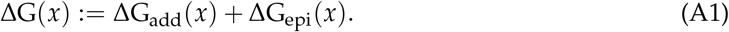

To specify each of the terms ΔG_add_(*x*) and ΔG_epi_(*x*), let *β*_{*k*},*α*_ be the additive contribution to the free energy of folding of having allele *α* at position *k*, and *β*_{*k*_1_,*k*_2_},*α*_1_*α*_2__ be the contribution of the interaction between allele *α*_1_ at site *k*_1_ and allele at site *k*_2_ in the pairwise interaction model. Then

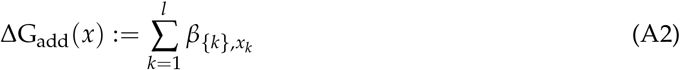

is the total additive contribution to the folding energy. For the pairwise interaction model we let

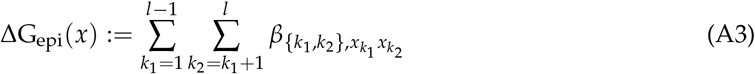

be the epistatic contribution to the free energy of folding. For the independent epistasis model we instead let ΔG_epi_(*x*) be an independent random draw from a normal distribution with mean 0 and variance 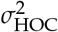.

It now remains to specify a procedure for assigning values to the *β*_{*k*},*α*_ and *β*_{*k*_1_,*k*_2_},*α*_1_*α*_2__. In drawing these coefficients, we need to ensure that ΔG_epi_(*x*) is a pure epistatic contribution, that is, that the average epistatic effect of any given point mutation over sequence space is zero, and would also like to set the average energy over all of sequence space to *lμ*_add_. To do this, we draw these coefficients from multivariate normal distributions with covariance matrices chosen to enforce the necessary constraints.

In particular, for each site *k*, we choose the *β*_{*k*},*α*_ from an *a*-dimensional normal distribution that has mean vector (*μ*_add_,…, *μ*_add_) and covariance

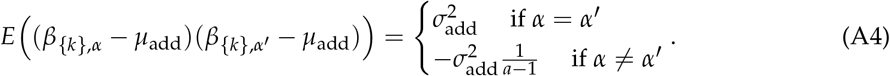

These coefficients are drawn independently for each site *k*, and the marginal distribution for each *β*_{*k*},*α*_ is a normal distribution with mean *μ*_add_ and variance 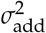 as required.

Turning to the epistatic component of the free energy of folding for the pairwise interactions model, for each pair of sites {*k*_1_, *k*_2_} with *k*_1_ < *k*_2_, we draw the *β*_{*k*_1_,*k*_2_},*α*_1_*β*_2__ from an *a*^2^-dimensional multivariate normal distribution with mean (0,…,0) and covariance

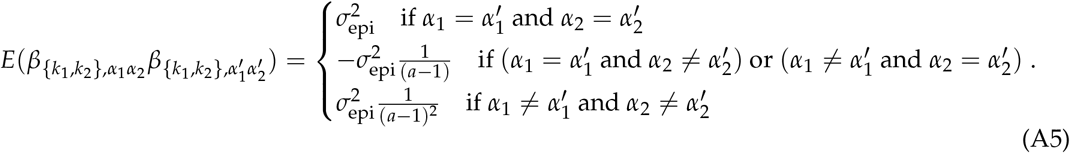

These coefficients are then drawn independently for each such pair of sites {*k*_1_, *k*_2_}. Furthermore, the marginal distribution for each *β*_{*k*_1_,*k*_2_},*α*_1_*α*_2__ is normal with mean 0 and variance 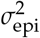 as required.

### Appendix A.2 Properties of the *β*{_*k*}α_ and *β*_{*k*_1_,*k*_2_},*α*_1_*α*_2__

As mentioned above, the covariance matrices for *β*_{*k*}*α*_ and *β*_{*k*_1_,*k*_2_},*α*_1_*α*_2__ are chosen to enforce two sets of constraints. First, for each *k*

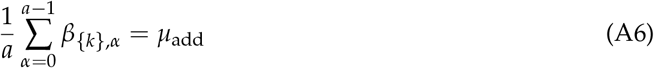

must be satisfied, so that the average contribution to the free energy of folding across all the alleles at any given site is *μ*_add_. This implies that the mean additive folding energy over all possible sequences is *lμ*_add_, i.e. 〈ΔG_add_(*x*)〉*x lμ*_add_.

Second, for each pair of sites {*k*_1_, *k*_2_}, we specify that

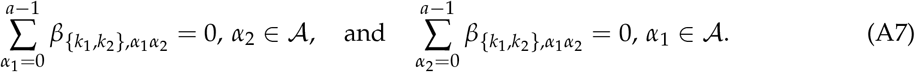

This second set of constraints ensures that the average epistatic effect of any given point mutation over sequence space is zero, i.e. that ΔG_epi_ is a pure epistatic term with no additive component. It also follows that the mean epistatic energy over all possible sequences is 0, i.e. 〈ΔG_epi_(*x*)〉_*x*_ = 0.

To see that constraint (A6) is satisfied, we calculate

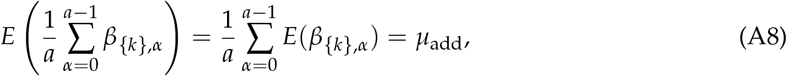

and using (A4),

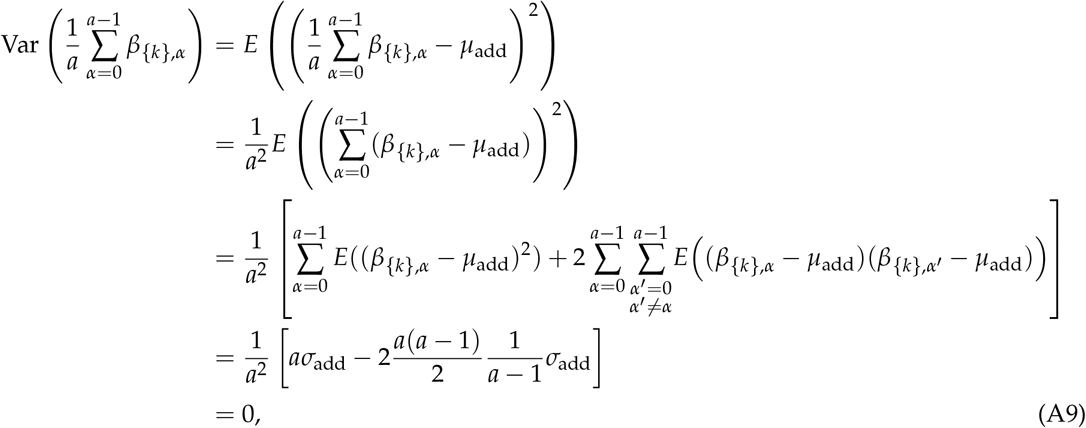

so that constraint (A6) is satisfied with probability 1.

Similarly, to see that constraint (A7) is satisfied, we calculate

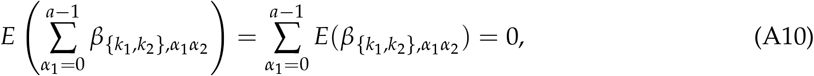

and using (A5),

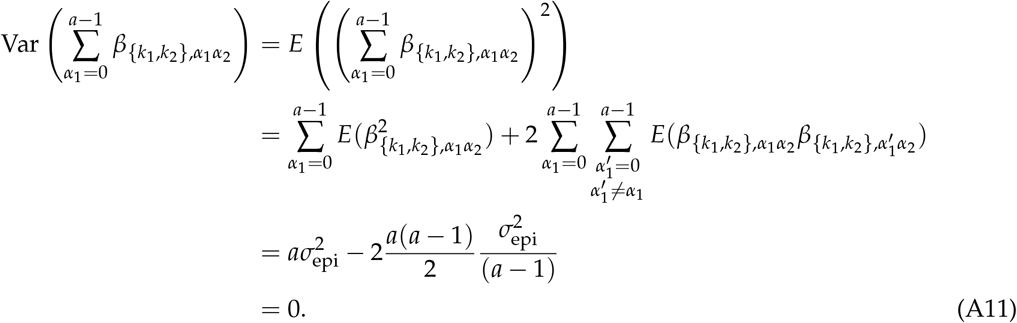

It follows from (A10) and (A11) that the first set of constraints in (A7) is satisfied with probability 1. By symmetry between *α*_1_ and *α*_2_, the same follows for the second set of constraints as well.

## Appendix B

### Appendix B.1 Expected variance of epistatic energy of distance classes under the random field model

In this section, we derive an analytical formula for the expectation of the realized variance in folding energy due to epistasis at a given distance from a focal genotype. Figure 3 was generated using this result. We arbitrarily fix a wild type sequence wt and look at all sequences that are at fixed Hamming distance *d* from this wild type, where the Hamming distance **d**(*x, x*′) between two sequences *x* and *x*′ is defined as the number of sites where the two sequences differ. Let 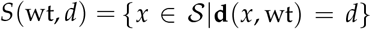 denote the set of sequences at distance *d* from the wild type wt. For some random function *f* defined on sequence space 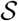, e.g. a random field model, we want to understand how variable the function will be among sequence at distance *d* from the wild-type. We quantify this variability in terms of the sample variance at distance *d*:

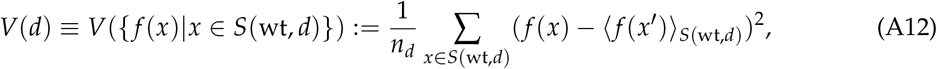

with 〈·〉_*T*_ denoting the mean taken over the set *T* and

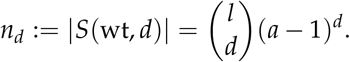

Because *V*(*d*) is itself a random variable and will take a different value for each realization of the random function, we quantify the typical behavior at distance *d* by the expected value of *V*(*d*), i.e. *E*(*ν*(*d*)). We will derive our results for *f* = ΔG_epi_, but they hold for any random field *f* that has the same covariance structure as ΔG_epi_ (i.e. isotropic, pure pairwise interactions).

We start by calculating the covariance function of

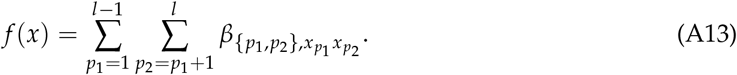

We obtain

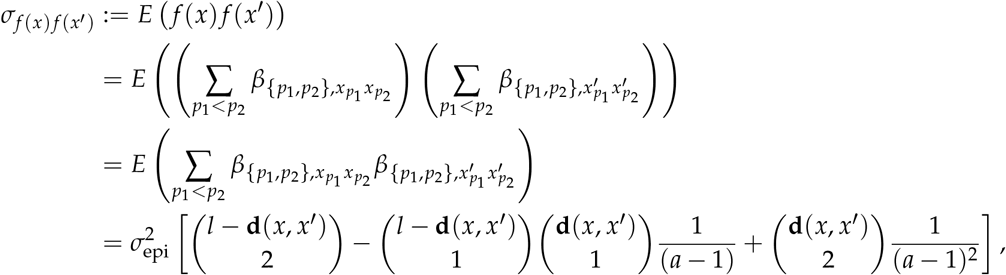

where first we used the fact that the *β*_{*p*_1_,*p*_2_}_ coefficients for different positions {*p*_1_, *p*_2_ } are independent, then broke up the sum according to position pairs at which *x* and *x*′ agree in 0,1, or 2 sites, and used the formula for the covariance between the coefficients *β*_{*p*_1_,*p*_2_},*α*_1_ *α*_2__ given in (A5). We note that *σ*_*f*(*x*)*f*(*x*′)_ is a function of **d**(*x, x*′), and we will use the notation

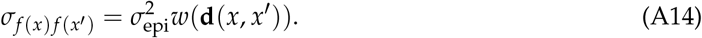

Now, turning to calculating the expected sample variance, we apply the well-known formula for possibly correlated random variables [51]

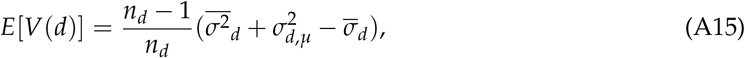

where the three quantities:

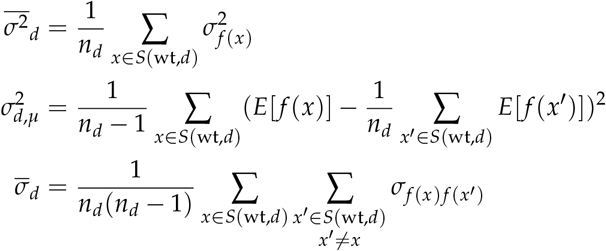

are the mean of the variances, the variance of the means, and the mean covariances, respectively. We now derive each of these three quantities individually.

First, we address the mean of variances 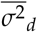. Since, by (A14), we have 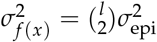 for all 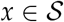, we also have

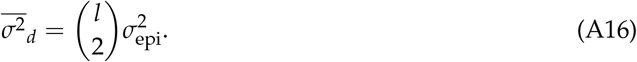

Next, for the variance of the mean energy 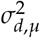, since the mean energy *E*[*f*(*x*)] is constant across all 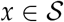, we have

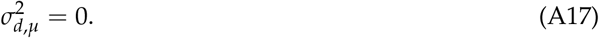

Finally, for the mean covariance 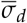, since the covariance *σ*_*f*(*x*)*f*(*x*′)_ only depends on **d**(*x, x*′) by (A14), the second sum in the definition of 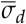 is the same for all *x*, thus we can arbitrarily fix a sequence *x* and write

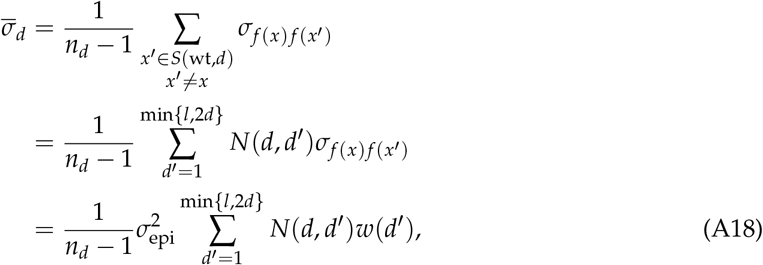

where *x*′ is an arbitrary sequence at distance *d* to wt and at distance *d*′ to *x*,

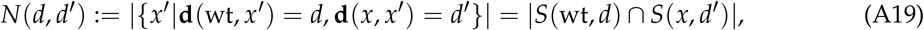

is the number of such sequences, and we used (A14) in the last equation. Note that

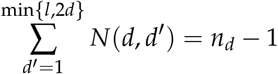

is the total number of sequences in *S*(wt, *d*) minus the focal sequence *x*.

Our final task is to count *N*(*d, d*′). First we pick *d* ≥ *s* ≥ 0 sites out of *d* sites on which *x* and wt differ and set the states of *x*′ on these sites to be the same as wt. The number of choices: 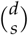. Second, since **d**(*x*′, wt) = *d*, we must choose *s* sites out of the *l* − *d* sites where *x* and wt are identical and set them to be one of the *a* − 1 states for *x*′. The number of choices is 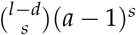. Third, now we have *x* and *x*′ differ on 2s sites and since **d**(*x, x*′) = *d*′, we need to choose *d*′ − 2*s* sites for *x*′ out of the *d* − *s* sites whose states we have not decided yet and set the states of *x*′ to be one of the *a* − 2 states that is different from both *x* and wt. The number of choices is 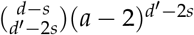

Putting these together, we obtain

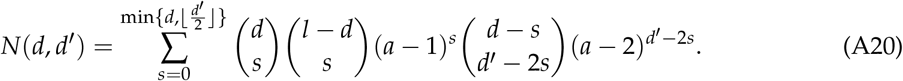

Returning to (A15), equations (A16), (A17), and (A18) yield

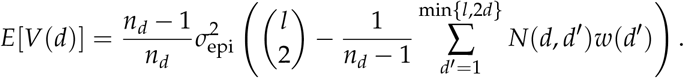

## Appendix C

### Appendix C.1 Bivariate normal approximation

We approximate (ΔG_add_, ΔG_epi_) with a bivariate normal distribution with mean vector

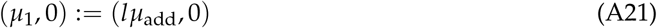

and covariance matrix

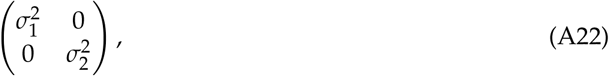

where 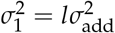 and 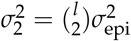 for the pairwise epistasis model and 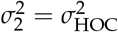 for the independent epistasis model. Thus, in this approximation, the total folding energy ΔG = ΔG_add_ + ΔG_epi_ is also normally distributed, with mean *μ* := *μ*_1_ and variance 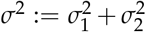. Using this normal approximation, we give an analytical justification for the phenomenon observed in Figure1g, that although the effect of epistasis is small, it is nonetheless crucial for folding. We shall use two quantities to measure the strength of this phenomenon. For the smallness of the epistatic effect, we use the measure 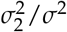, the fraction of the energy variance across all sequences accounted for by the variance of the epistatic energy. For the importance of the epistatic effect, we use the measure *E*(Δ*G*_add_|Δ*G* < 0), the mean additive energy of viable sequences. If this mean is far above the viability threshold 0, it indicates that, on average, epistasis must make a substantial contribution to the ability of viable sequences to fold.

We now analytically approximate the conditional expectation *E*(Δ*G*_add_|Δ*G* < 0). We use a classical result that is referred to as the regression towards the mean formula for a pair of normally distributed random variables. If (*X, ϒ*) has normal distribution with mean (*μ_X_, μ_ϒ_*) and covariance matrix

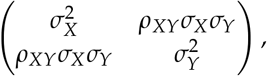

then the regression towards the mean formula describes how the means change if we condition on one of the variables being below some cutoff value *c*:

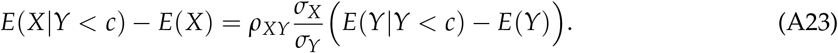

Applying this formula to Δ*G* and Δ*G*_add_, and the condition that a sequence is viable, i.e. Δ*G* < 0, we obtain

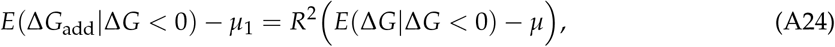

where

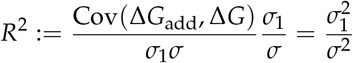

is calculated using the fact that Δ*G* = Δ*G*_add_ + Δ*G*_epi_ and that the additive and epistatic energies are uncorrelated by (A22). Also by (A22), *μ* = *μ*_1_, hence we can express the mean additive folding energy of viable sequences from (A24) as

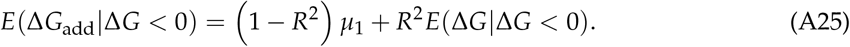

The conditional mean on the right hand side of the equation above can be calculated as

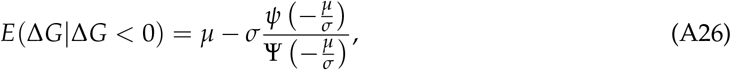

where *ψ* and Ψ are the PDF and CDF of the standard normal distribution, respectively. As *μ* becomes large compared to *σ, E*(Δ*G*|Δ*G* < 0) approaches 0, therefore, returning to (A25), we obtain the estimate

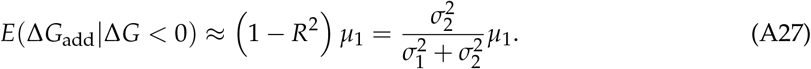

This means that no matter how small the epistatic effect is, measured by 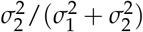, if the mean of the additive energy *μ*_1_ is large enough in comparison, the role of the epistatic energy in protein folding is crucial.

Plugging in our model parameters as given in (A21) and (A22), and using the approximation *l* − 1 ≈ *l*, the estimate in (A27) becomes

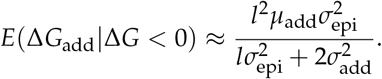

The choice of parameters *μ*_add_ = 1, 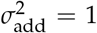, and 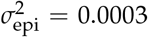 used in our simulations then yield *E*(Δ*G*_add_|Δ*G* < 0) ≈ 22.6, which is very close to 22.5, the mean additive energy of sequences observed in the simulation.

